# Competition between IAV subtypes through heterosubtypic immunity modulates re-infection and antibody dynamics in the mallard reservoir

**DOI:** 10.1101/063933

**Authors:** Neus Latorre-Margalef, Justin D. Brown, Alinde Fojtik, Rebecca L. Poulson, Deborah Carter, Monique Franca, David E. Stallknecht

**Affiliations:** Southeastern Cooperative Wildlife Disease Study, College of Veterinary Medicine, Department of Population Health, The University of Georgia, 589 D. W. Brooks Drive, Athens, Georgia 30602, USA; Department of Biology, Lund University, Ecology Building, 223 62 Lund, Sweden; Pennsylvania Game Commission, Pennsylvania State University, Animal Diagnostic Laboratory, Orchard Road, University Park, Pennsylvania 16802, USA; Poultry Diagnostic and Research Center, College of Veterinary Medicine, Department of Population Health, The University of Georgia, 953 College Station Road, Athens, GA 30602, USA

## Abstract

Our overall hypothesis is that host population immunity directed at multiple antigens will influence the prevalence, diversity and evolution of influenza A virus (IAV) in avian populations where the vast subtype diversity is maintained. To investigate how initial infection influences the outcome of later infections with homologous or heterologous IAV subtypes and how viruses interact through host immune responses; we carried out experimental infections in mallard ducks (*Anas platyrhynchos*). Mallards were pre-challenged with an H3N8 low-pathogenic IAV and were divided into six groups. At five weeks post H3N8 inoculation, each group was challenged with a different IAV subtype or the same H3N8. Two additional pre-challenged groups were inoculated with the homologous H3N8 virus at weeks 11 and 15 after pre-challenge to evaluate the duration of protection, which showed that mallards were still resistant to re-infection after 15 weeks. There was a significant reduction in shedding for all pre-challenged groups compared to controls and the outcome of the heterologous challenges varied according to hemagglutinin (HA) phylogenetic relatedness between the viruses used. There was a boost in the H3 antibody titer after re-infection with H4N5, which is consistent with original antigenic sin or antigenic seniority and suggest a putative strategy of virus evasion. These results imply strong competition between related subtypes that could regulate IAV subtype population dynamics in nature. Collectively, we provide new insights into within-host IAV complex interactions as drivers of IAV antigenic diversity that could allow the circulation of multiple subtypes in wild ducks.

**Author summary:** Many features of pathogen diversification remain poorly explored although host immunity is recognized as a major driver of pathogen evolution. Influenza A viruses (IAVs) can infect many avian and mammalian hosts, but while few IAV subtypes circulate in human populations, subtype diversity is extensive in wild bird populations. How do these subtypes coexist in wild avian populations and do they compete within these natural host populations? Here we experimentally challenged mallard ducks with different IAVs to study how an initial infection with H3N8 determines the outcome of later infections (duration of infection and virus load) and antibody responses. There was complete protection to re-infection with the same H3N8 virus based on virus isolation. In addition, there was partial protection induced by H3N8 pre-challenge to other subtypes and development of heterosubtypic immunity indicated by shorter infections and reduction in viral load compared to controls. This indicates that subtype dynamics in the host population are not independent. Amongst H3N8 pre-challenged groups, the highest protection was conferred to the H4N5 subtype which was most genetically related to H3N8. The H4N5 challenge also induced an increase in H3 antibody levels and evidence for antigenic seniority. Thus, previous infections with IAV can influence the outcome of subsequent infection with different IAV subtypes. Results not only have relevance to understanding naturally occurring subtype diversity in wild avian populations but also in understanding potential outcomes associated with introduction of novel viruses such as highly pathogenic IAV H5 viruses in wild bird populations.

**Author contributions:** Conceived and designed the experiments: NLM, DES. Performed the experiments: NLM, JDB, AF, DC, MF, DES. Contributed reagents/materials/analysis tools: NLM, JB, AF, RLP, DES. Analyzed the data: NLM, DES. Wrote the paper: NLM, JDB, AF, RLP, DC, MF, DES

## Introduction

Diversification is a common feature in pathogen populations and this often involves the evolution of antigenic variants (1). Examples of this exist in various pathogen systems: viruses (influenza A virus, Dengue virus, Bluetongue virus, and rotaviruses), bacteria (*Borrelia spp, Neisseria meningitis, Pneumococcus*) and protozoan parasites (*Plasmodium spp.* and trypanosomes). Antigenic variation within specific hemagglutinin (HA) and neuraminidase (NA) subtypes is well described with influenza A viruses (IAV) as this is an important consideration in developing vaccines and vaccination strategies associated with seasonal influenza viruses in humans and IAV affecting domestic livestock and poultry. Though multiple IAV subtypes circulate in these host systems, antigenic interactions between these subtypes are equally important but less understood. Shared epitopes between different HA subtypes associated with the HA stalk have been described and these may be important target epitopes for universal IAV vaccines. Immunity to these shared epitopes also may provide partial protection in subsequent infections with heterologous IAV and could potentially influence clinical outcome and regulate subtype diversity in host populations through competition (2).

IAV have the capacity to infect many different hosts, from birds to mammals, however the vast majority of influenza subtype diversity is found in wild birds, especially waterfowl, gull, and shorebird populations where low-pathogenic IAV (LPIAV) representing 16 HA and 9 NA subtypes are maintained (3). In these wild bird populations, IAV subtypes can be represented in many HA/NA combinations. Many subtypes co-circulate at the same time and same location (4–6). In addition, individual birds are often co-infected or sequentially infected with multiple IAV subtypes in a given season or year (7, 8)

Experimental infections have demonstrated that initial infection with a specific IAV induces immune responses and protection against infection with homologous strains (9). Additionally, initial infections with a virus could induce partial protection to heterologous subtypes in experimental settings (9–12) and several studies have shown that LPIAV pre-exposure protects against a lethal highly pathogenic IAV (HPIAV) challenge (10, 13, 14). Naturally infected mallards (*Anas platyrhynchos*) in the wild also exhibit patterns of re-infections indicating heterosubtypic cross-immunity (8). Despite these observations little is known about the mechanisms, extent, and persistence of these responses and the outcomes of re-exposures that have critical significance to understanding the maintenance of IAV antigenic diversity.

In this study, mallards were pre-challenged with a strain of H3N8 LPIAV, a common subtype in waterfowl, and pre-challenged ducks were subsequently re-challenged with the same strain (homologous challenge) or with different strains representing various subtypes (heterologous challenges). Groups of ducks in the homologous challenge were exposed at different time intervals to evaluate long-term antibody responses and potential protection. The different strains used in the re-challenge represented a gradient in the degree of phylogenetic relatedness between the HA.

The objective of the present study was to experimentally mimic re-infections that commonly occur in mallards and other ducks in nature to determine the effects on susceptibility, duration and intensity of viral shedding and to characterize the humoral immune response.

## Results

### Variation in Virus Load and Duration of shedding

#### Homologous challenge

Individuals were initially inoculated at 4 week of age of with LPIAV H3N8 (pre-challenge) and all were susceptible to infection and shed virus. Study design can be found as supporting information (S1 Table). Virus replication occurred in all the control groups for the H3N8 re-challenge where Ct-values from the matrix RRT-PCR were lowest in cloacal (CL) samples at 2 dpi, indicating a peak in virus load/shedding; Ct-values increased over time (Fig 1). None of the swabs from H3 pre-challenged groups were positive by virus isolation after the homologous re-challenge; however some swabs tested positive by matrix RRT-PCR (Fig 1-12, 3, and 4 positive samples in the 5, 11, and 15 week re-challenge groups, respectively). In all cases (5-, 11-, and 15-week re-challenge), Ct-values were higher in pre-challenged birds than in controls indicating that pre-challenged birds shed fewer virus copies than naïve birds when challenged with the same strain. The variation in Ct-Values between groups was evaluated using linear mixed models as described later in the methods section. The best models (S2 and S4 Tables) for the groups 5 and 15 weeks post-challenge included dpi, group and the interaction indicating that the rate of clearance was different for the different groups (a different slope per group). For the group re-challenged after 11 weeks the best model included dpi and group (S3 Table). In all groups oropharyngeal (OP) shedding was generally lower than CL shedding (S1 Fig). Individuals did not show symptoms of disease and all gained weigh throughout the trial.

**Fig 1.**
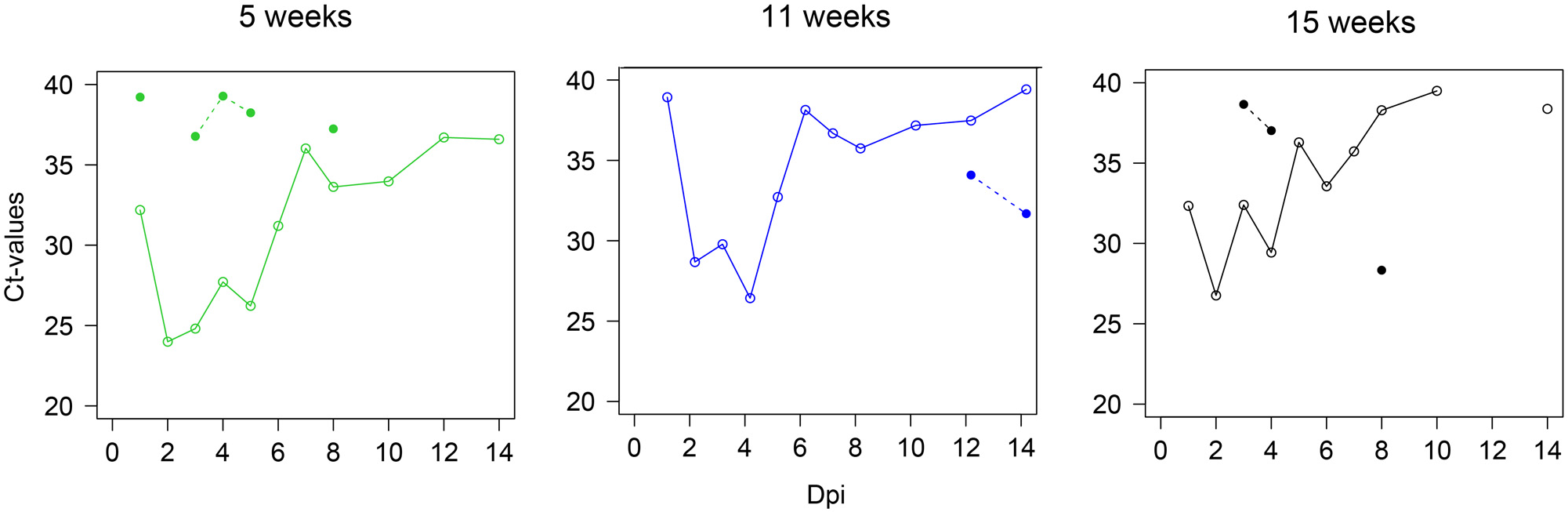
Pre-challenge reduces viral shedding in homologous challenge. Patterns of CL shedding in pre-challenged and control groups for different age and time intervals at re-challenge. The panels show the variation in Ct-values (mean Ct-values per group, no data points indicate negative RRT-PCR, lines are drawn between days with consecutive positive results) for the different groups: 5 weeks, 11 weeks, 15 weeks intervals between H3N8 homologous infections. Continuous lines and circles indicate the control groups and dashed lines with filled dots indicate the H3N8 pre-challenged groups. This means that pre-challenge induced protection and suppresses CL shedding.

Based on comparisons of viral shedding of H3N8 during primary challenge (H3 pre-challenge and control groups) younger individuals (4 and 9 weeks of age) had lower Ct-values (higher viral shedding) than older individuals (Fig 2) and duration of infection was shorter for older birds. Based on the best model by AICc, there were significant differences in Ct-values based on dpi and age group (S5 Table). The mean duration of infection for H3N8 in controls was 4.8 to 8.4 days (primary infections) (Fig 3) and there was a significant difference reduction in the duration of infection in older birds (p-value = 0.02).

**Fig 2.**
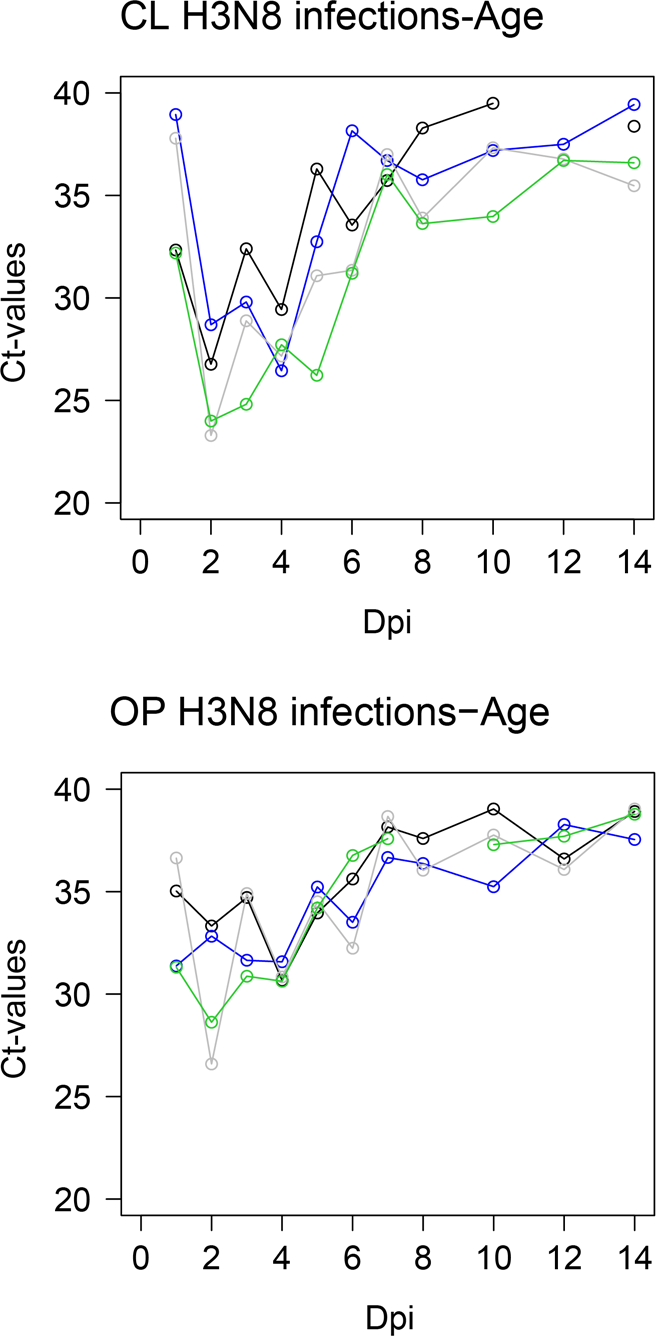
H3N8 CL viral shedding for the different age groups. Graph shows the variation in Ct-values for the different age groups: grey lines are 4 weeks of age, green are 9 weeks, blue are 15 weeks and black are ducks at 19 weeks of age. A) Cloacal and b) Oropharyngeal shedding.

**Fig 3.**
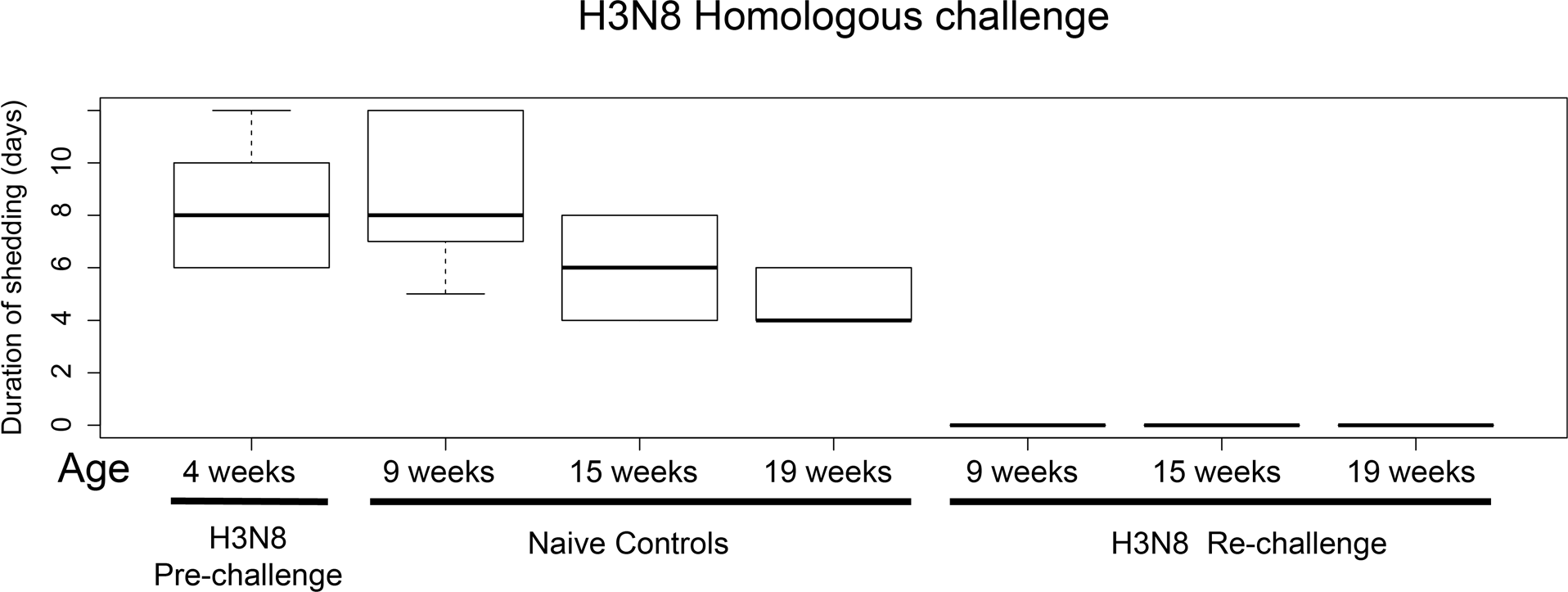
Duration of shedding for H3N8 homologous challenge groups and controls. Pre-exposure protects against homologous re-challenge in all groups based on virus isolation from CL samples. Duration of infection is reduced in older age groups.

#### Heterologous challenge: subtype interaction

All birds in the H3N8 pre-challenge were susceptible to infection and all were negative by virus isolation on CL swabs on 21 and 35 dpi, prior to transfer to poultry isolators for the re-challenge trial. All viruses replicated in control birds as detected by virus isolation and RRT-PCR, except for the H12N5 strain (therefore data from the H12N5 groups could not be used for comparison, results for this group are shown in S2 Fig). For the H4N5, H10N7 and H6N2 viruses, Ct-values in H3 pre-challenged groups were higher than in the control groups indicating a reduction in virus replication and partial protection compared to controls (Fig 4). Ct-values from naïve (control) and pre-challenged ducks were evaluated using mixed models. There were significant differences in Ct-values between the pre-challenged group H3N8 x H4N5 group and the H4N5 controls and best models included dpi, group and the interaction between dpi and group, which means a difference in clearance rate for the different groups (S6 Table). For the H3N8 x H10N7 and H3N8 x H6N2 groups, the best models indicated significant differences in Ct-values when compared with their respective controls and both dpi and group were significant (S7-S8 Tables). Summaries of the model selection (AICc) and significance values are provided as supplementary material (S6-S8 Tables). If differences in Ct-values from pre-challenged and controls are translated into “log” (as a difference of 3.3 Ct-values is equivalent to one log) the reduction in shedding ranges between 0.7 and 4.75 logs depending on the groups.

**Fig 4.**
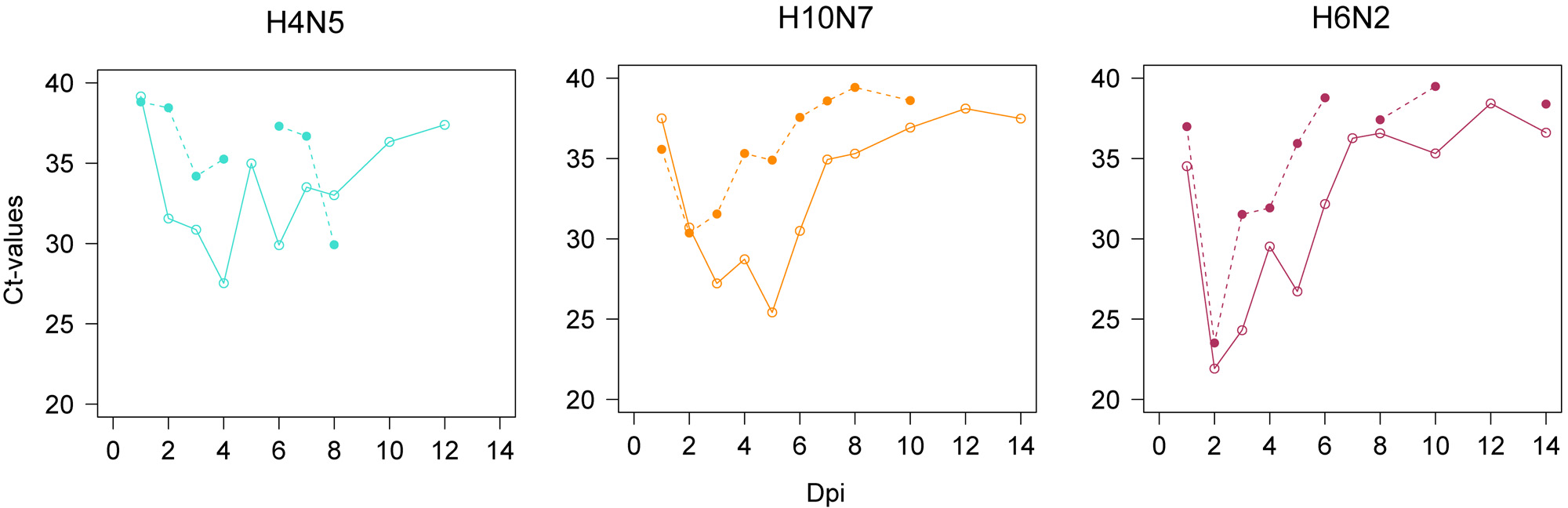
Pre-challenge reduces cloacal (CL) viral shedding after heterologous challenge. Birds were pre-challenged (dashed lines) or not (solid lines) with H3N8 and challenged with different LPIAV viruses/strains. CL shedding showing the variation in Ct-values for the different heterologous challenge groups: H4N5, H10N7, H6N2. Continuous lines and circles indicate the control groups and discontinuous lines indicate the H3N8 pre-challenged groups.

Mean duration of infection (as determined by detectable viral shedding) in the control groups at 9 weeks of age was 8 days (sd 2.90) (range 7 - 8.8 days) while in the pre-challenged groups the mean duration of shedding was 2.6 days (sd 2.75) ranging from 0 days in the homologous challenge to 2.4 days for H3N8 x H4N5 group with only two individual birds infected (two CL samples), to 3.6 days for the H3 x H10 group where all individuals were infected (9 CL isolates recovered), and 5 days for H3N8 x H6N2 group where individuals were also infected (9 CL isolates). There were no differences in the duration of shedding between the different subtypes (p-value = 0.83) on the naïve control groups. There were significant differences in the duration of infection between pre-challenged groups (p-value = 0.034) (Fig 5).

**Fig 5.**
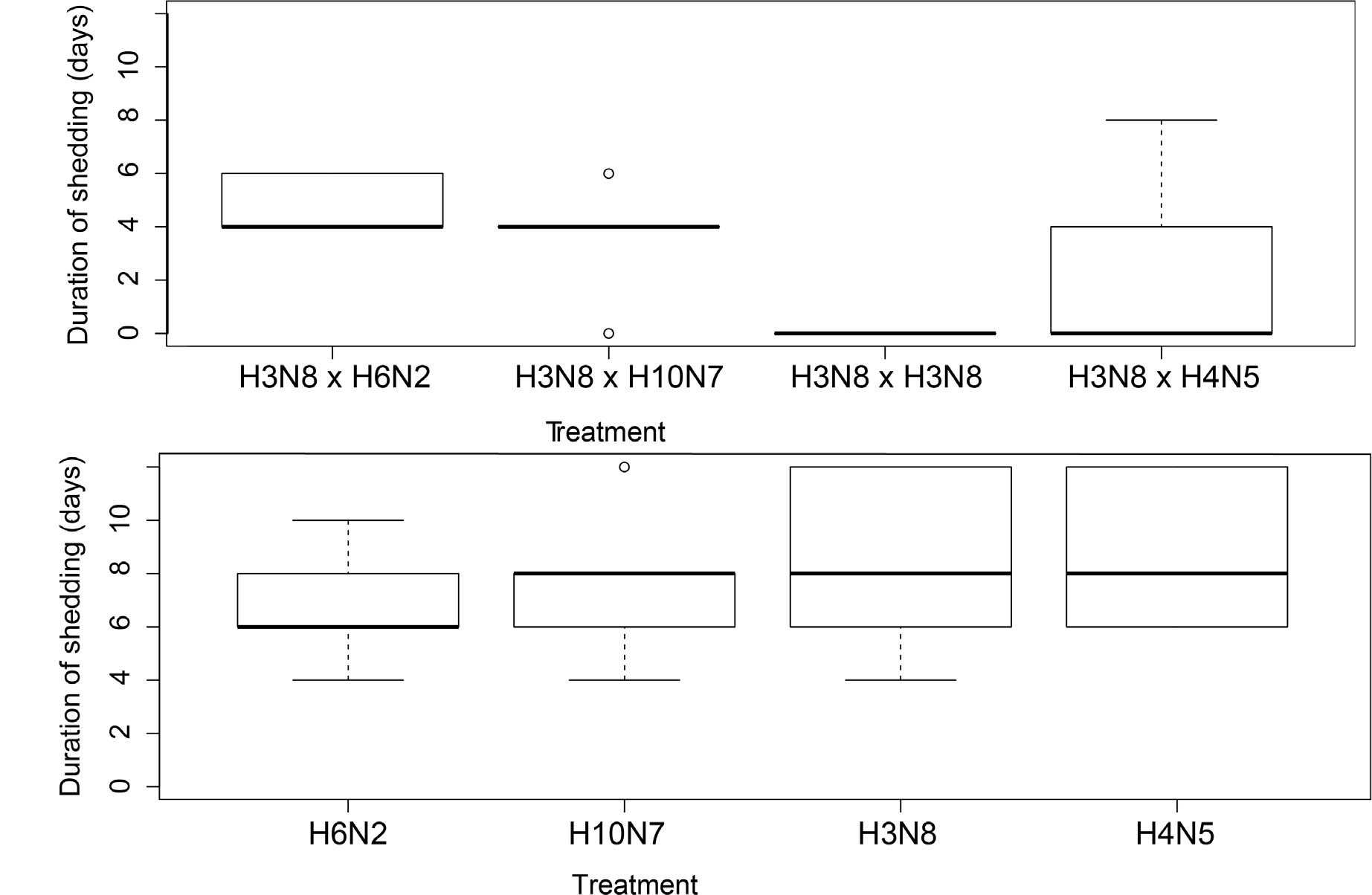
Pre-exposure reduces the duration of shedding. Duration of infection for viruses used in heterologous challenge (based on virus isolation for CL samples): A) controls and B) H3N8 pre-challenged groups.

To test if genetic distance between HA subtypes was an important determinant in the strength of protection the relationship between HA amino acid sequences and reduction of infection was evaluated. There was a negative correlation between the amino acid distance and the reduction in duration of infection per group (R^2^= 0.94, ranging from 100% to 29% depending on the subtype in the re-challenge) (Fig 6).

**Fig 6.**
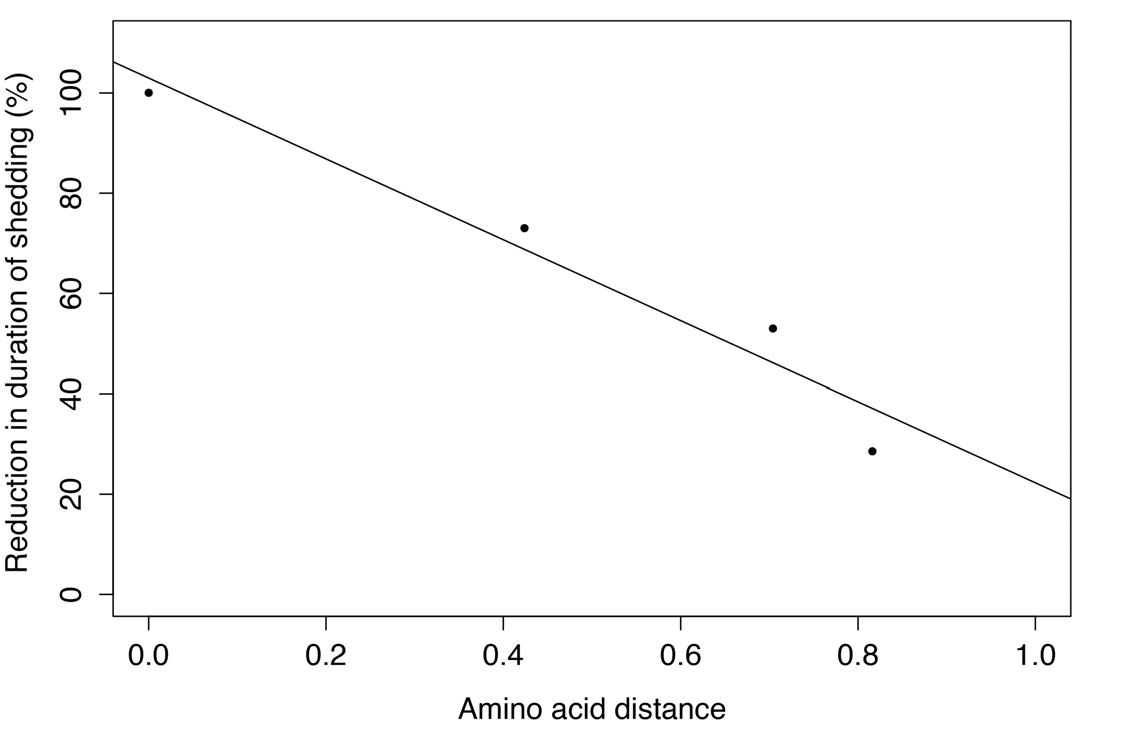
Correlation between amino acid distance between HA in infections and the reduction in the duration of infection (days).

### Infectivity: Variation in Ct-values and virus isolation

We evaluated the factors that influenced isolation success of RRT-PCR positive samples and found that there was a significant negative relationship between isolation and Ct-values. As anticipated, samples with high Ct-values tended to have a lower isolation success that those with low Ct-values (high number of viral RNA copies). Remarkably both the H3N8 pre-challenge treatment (whether samples derived from primary or secondary infections) and dpi were significant in the model. This indicates that the probability of detecting IAV in RRT-PCR positive samples by virus isolation varies related to infection history and the duration of infection. This was not dependent on sample type (CL or OP) (S9 Table).

### Serology

#### Long-term Persistence of antibodies in the Homologous challenge

All individuals were seronegative prior to infection and seroconverted as measured by NPELISA on 14 dpi after pre-challenge and all but one also had H3 specific antibodies as measured by MN using the homologous antigen. NP-antibodies and H3 specific antibodies were detectable 5 weeks after initial H3N8 infection. There was no boost in the H3 antibody response in the 5 week interval group (p-value = 0.326). For the 11 and 15 week challenge groups, there was a slight decrease in antibody titer over time with a boost bothin the NP and H3 specific antibody responses following re-challenge (11 weeks, p-value = 0.003; 15 weeks p-value = 0.032) (Fig 7, panels a) and c)).

**Fig 7.**
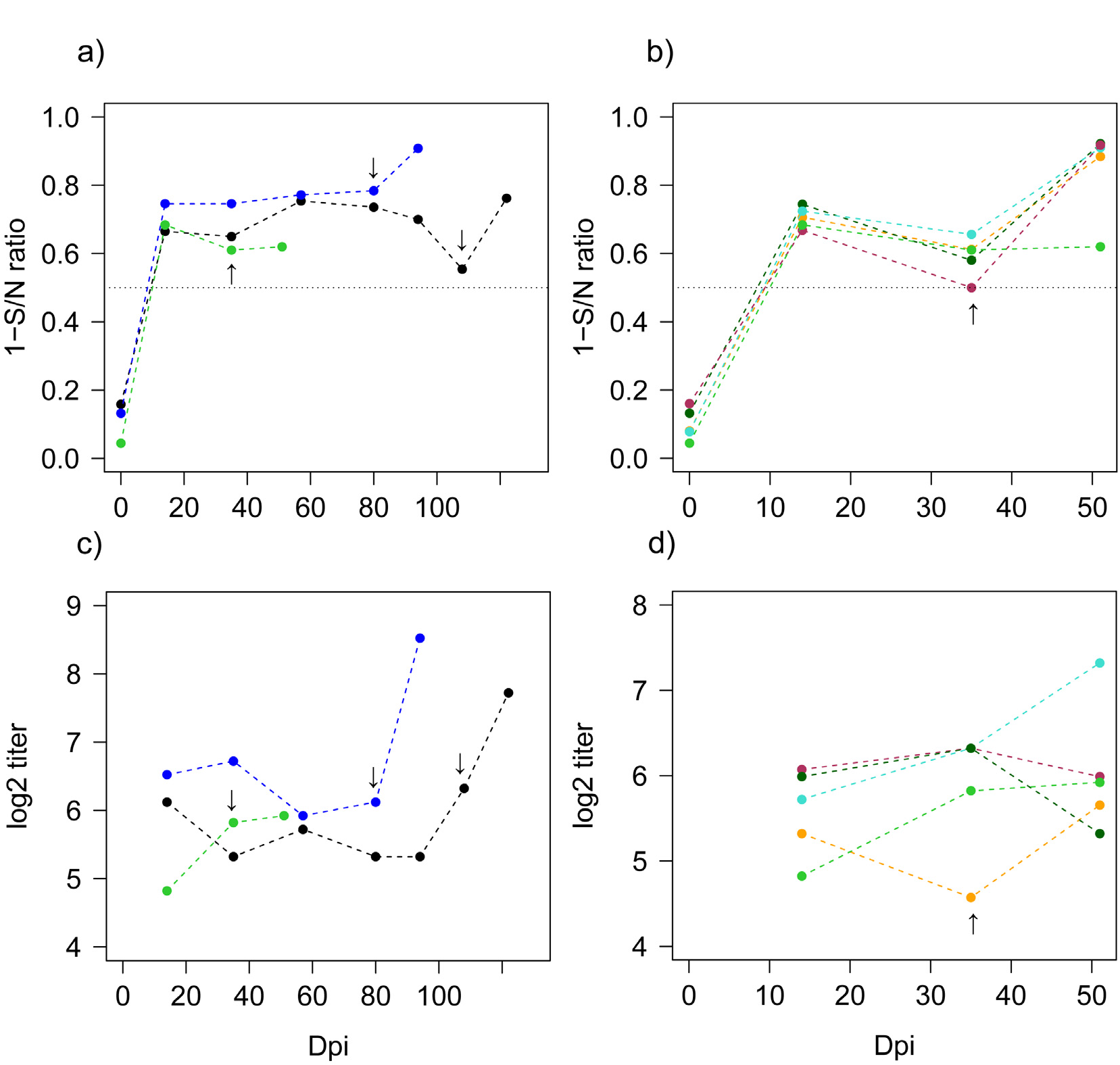
Antibody dynamics for the H3N8 homologous (panels a and c) and heterologous challenges (panels b and d). Data points indicate the average per group over time. Arrows indicate the timing of re-challenge for the different groups. The different colors indicate groups pre-challenged with H3N8 (green is the group re-challenged after a 5 weeks interval, blue are 11 weeks and black are at 15 weeks). The different colors indicate the different challenge groups: the groups challenged with H4N5 are light blue, H10N7 in orange, H6N2 in red, H12N5 in dark green and H3N8 (homologous challenge for reference) are green. Continuous lines are for controls and dashed for H3N8 pre-challenged., a) and b) NP-ELISA ratios levels (expressed here as one minus the ratio) c) and d) H3 response by MN (log_2_ transformed titers) over time.

#### Antibody responses to heterologous challenge

All individuals tested seronegative by NP-ELISA prior to infection. Individuals pre-challenged with H3N8 seroconverted on both NP-ELISA and H3 MN by 14 dpi. Based on the NP-ELISA response there was an increase in S/N ratios in all groups after the secondary challenge including the group challenged with H12N5 (Fig 7, panel b). There were no differences in H3 MN titers between groups of birds assigned to the different heterologous groups after initial H3N8 infection (p= 0.261, 14 dpi) and prior to secondary infection (p= 0.242, 35 dpi). An increase in the H3 MN titer was observed in the H3N8 x H4N5 challenge group (p-value = 0.200) (Fig 7, S9 Table); antibodies to H4 were also detected (Fig 8). Similarly the group H3N8 x H10N7 showed a slight increase in the H3 MN titer. To evaluate the variation in H3 MN titers between H3 pre-challenged groups and the influence of time (i.e. prior to and after re-challenge) we used mixed models. The best model included the effect of dpi (S10 Table). All groups and most individuals developed detectable antibodies against the challenge viruses used in secondary challenges (Fig 7). MN results from sera collected at the termination of the experiment using a panel of H1-H15 prototype strains are shown in Figure 8. Most individuals had antibodies specific to the HA subtypes they were challenged with however, one sample MN positive to H1N1 was observed in the homologous group (5 weeks interval).

**Fig 8.**
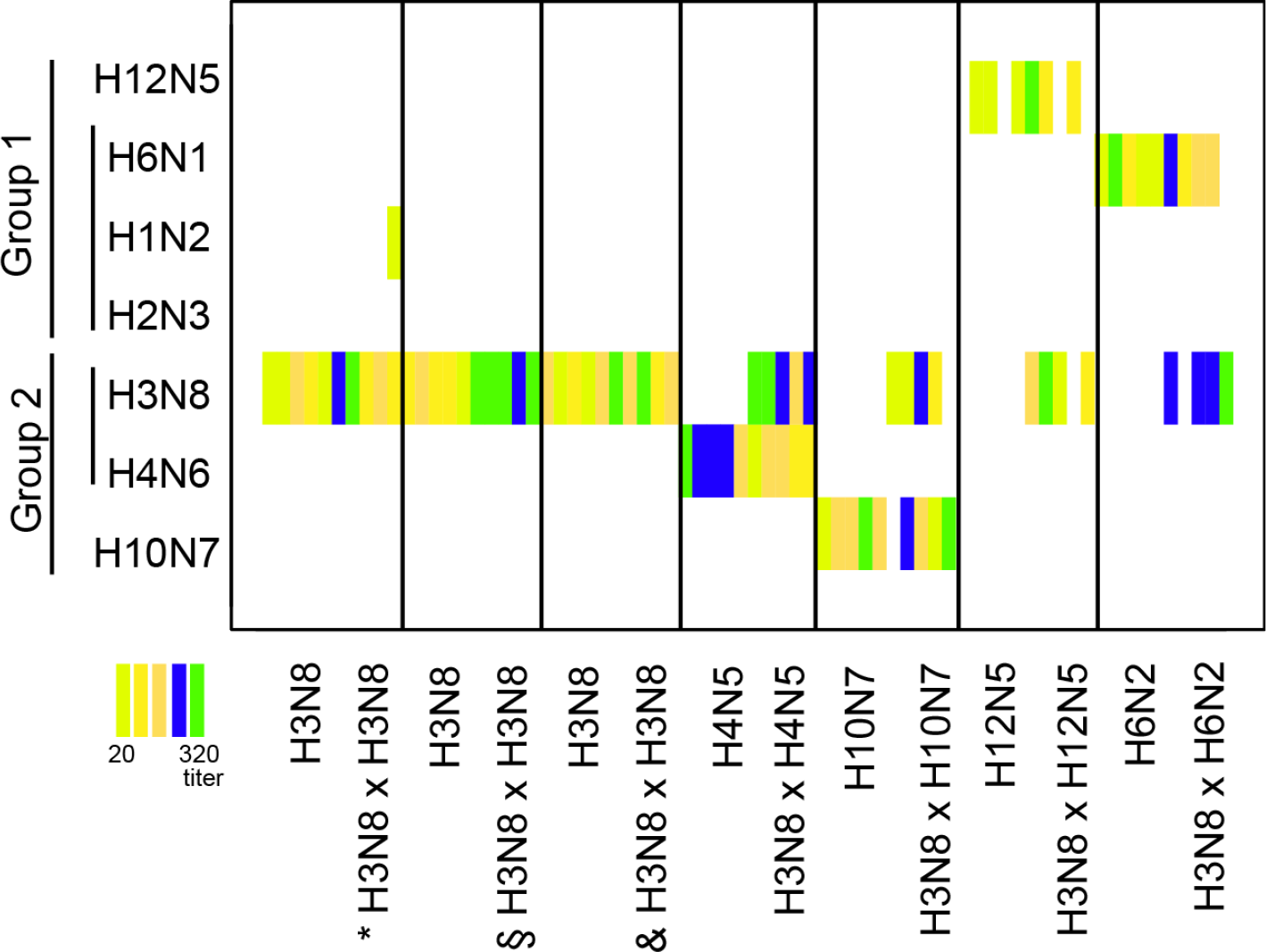
Heat map indicating virus neutralization profiles after secondary infection against a complete panel of H1-H12 and H14-H15 antigens for the different treatments (only antigens with positive MN included). Colored boxes indicate the titers based on the legend. The challenge groups are indicated on the horizontal axis while the vertical axis indicates the antigens used in the neutralizations and their phylogenetic relationships. The homologous challenge 5 weeks interval is indicated by a *, 11 weeks by § and 15 weeks interval by &.

## Discussion

Many IAV HA/NA subtype combinations coexist in duck, gull and shorebird populations and it is notable that their abundance and relative diversity varies over time and space (4–6). What determines whether a particular virus is more common than others? Is one subtype driving the abundance of other subtypes? How are subtypes interacting in the mallard-IAV system? Pathogen fitness or success is usually measured by transmission risk that could be based on different transmission parameters such as the duration of infection and pathogen load (15) as well as how long viruses could remain infectious in the environment (16, 17). Another important parameter is the development of protective immune responses characterized by the strength, specificity, duration, and potential response to subsequent exposures (25). Protective responses may be especially important in multi-strain/subtype pathogen systems such as occurs with IAV and ducks.

### Homologous challenge

Here, we studied the persistence of protection to homologous IAV challenge by initially infecting mallards with H3N8 and re-challenging three groups with the same virus at different time intervals. The initial H3N8 induced long-term protection against homologous re-infection for up to 15 weeks post-challenge which is much longer than expected. None of the re-challenged birds, regardless of time interval, shed virus as determined by virus isolation and that was true for all age groups and time intervals. Although some RRT-PCR positives were detected in re-challenged individuals, the Ct-values were significantly higher than in control groups indicating a low number of RNA copies possibly associated with non-infective virus. The observed protection and detectable antibody response is inconsistent with results from previous studies that concluded that infection conferred no protection against re-infection and that detectable antibody responses in ducks were short-lived (18, 19). It was considered that antibody responses may not be detectable or protective due to the truncated structure of the IgY. A possible explanation of this inconsistency is that these truncated antibodies neutralize IAV but lack HI activity (20). Our results are in agreement with other studies in both ducks and gulls that reported partial to complete protection against re-infection depending on the viruses, time between infections, host species and age and detection method (9, 21, 22). The long-term persistence of antibodies after natural infection, artificial challenge or vaccination has been reported in captive birds up to 6 to 9 months post-exposure (23–25) though it is not know which parameters (i.e. HI or MN titers) are correlated to protection. When examining patterns in H3N8 infections for the different age classes in the naïve controls all individuals were susceptible and competent to infection. The duration of infection and viral load in controls varied according to age. This is indicative of a different degree of exploitation, consistent with earlier findings (26), and that could also influence subsequent transmission risk and subtype diversity in cases were subtypes display a seasonal pattern.

### Heterologous challenge

Results from IAV heterologous challenge experiments of ducks, geese and gulls have reported varying levels of partial protection upon re-challenge (11, 29, 30, 35). Such protection has also been reported from field-based studies (8, 27) and from experimental work where birds were challenged with HPIAV and effects could be measured by morbidity/mortality responses(10, 13). It has also been reported that the order in which IAV subtypes are used to challenge the host is important. For instance, H3 appears to be poorly immunogenic and less protective as a primary infection compared to H4 that induced a stronger protective response (12). The variation in reported protection from previous studies may be the result of asymmetric responses associated with strain or subtype specific variation that may be dependent on the virus causing primary infections (11, 12, 28).

We observed a partial protection to re-challenge in the individuals subsequently infected with heterologous subtypes indicating development of heterosubtypic cross-immunity by H3N8. When assessing the outcome of re-challenge there was a significantly lower viral load and shorter duration of infection in CL samples for all the groups pre-challenged with H3N8 compared to the naïve controls. This shorter period of detectable viral shedding is in agreement with estimates from the field (29, 30) and was observed for all re-challenged viruses.

The extent of partial protection, as measured by a decrease in duration of detectable viral shedding following re-infection was influenced by the genetic relatedness of the HA. This suggests immune mediated competition trough cross-reactive responses (31, 32) and this could be related to both acquired humoral and cell mediated mechanisms (33). Broadly neutralizing antibodies have been defined across HA groups (34) and within HA group (35, 36) that are targeting conserved epitopes in the stalk (37, 38). Stalk antibodies are boosted upon re-infection and is it therefore thought that cross-reactive anti-stalk as well as anti-NA antibodies (39) can diminish the severity of the disease in re-infections. Studies in humans have found consistent patterns of cross-immunity within HA group with increased severity of the disease in cohorts exposed with an HA of the opposite group during childhood when studying age-specific mortality caused by 1918 H1N1 and by HPIAV H5 and H7 (40, 41). It has been proposed that extinction of influenza strains in humans could be driven by population immunity by stalk and NA antibodies (39) through competitive exclusion between strains (2). Moreover, initial infections with a specific virus increases the probability of later infections by viruses from a different HA clade and group in wild mallards (8). Comparable processes are likely acting in the avian IAV system where lineage replacement with strains from different continents has been reported (42, 43). Theory predicts that antigenic variants tend to organize as discrete non-overlapping strains (1) in populations where cross-reactivity between viruses induces competition (i.e. case of H13 and H16) contrasting to the situation where cross-reactivity induces enhancement or facilitation, like Dengue, and variants may be antigenically clustered (44). Ducks after the heterologous re-challenge shed viruses even if they were able to clear infections rapidly. An implication of that is the potential for selective processes like viral escape and antigenic drift to act in the same way as leaky vaccines (45, 46). For instance, that in turn would be driving antigenic evolution as observed for H5N1 HPIAV (47) or similarly for seasonal H3N2 in humans (48). Indeed, the estimates of divergence for some HA subtype indicate that divergence is relatively recent (40).

### Infectivity

Isolation success from RRT-PCR positive samples from secondary infections (re-challenge), as expected, was correlated with Ct-value, but was significantly lower compared to isolation results from primary infections in controls. This discrepancy has implications when interpreting field data based on molecular diagnostics rather than virus isolation and may explain why isolation rates from PCR positive samples often vary. This relationship also corroborates findings from the field where samples from adults have a lower isolation success than samples from juveniles (49) and where RRT-PCR positive samples used in experimental trials have not infected ducks (50). Therefore we propose caution when using transformations of Ct-values (proxy for RNA copies) to EID_50_/ml or TCID_50_/ml by a standard curve based on a virus grown at optimal conditions such as cell culture or embryonated eggs as infectivity in hosts varies according to several parameters (age of host, previous exposure, specific virus and infection dose…).

### Antibody dynamics

Antibody levels (NP and H3 MN), after H3N8 infection, decreased over time but most individuals remained positive for the duration of the experiment (15 weeks); all birds remained protected against homologous challenge. There was a substantial boost in the antibody responses after homologous challenge for the long-term groups (11 and 15 weeks) but not for the group re-infected after 5 weeks where a rapid blocking of the infection may not have activated antibody recall. Additionally, the hyperimmune sera after homologous challenge did not cross-react with other IAV subtypes, except for one individual positive by H1 at a low titer of 20.

The heterologous re-infections resulted in a boost in the NP-antibody responses in all H3 pre-challenged groups. Interestingly most of the individuals had neutralizing antibodies against the HA antigens they had been exposed to. This also includes the H12N5 that replicated poorly but resulted in serological imprinting in three of the five individuals per group. Additionally no cross-reactivity to other subtypes was observed when tested by MN with a panel of HA (H1-H12 and H14-H15). H3-specific antibodies were detected by MN after H3N8 primary infection and persisted in the majority of individuals until the end of the experiment. There were interesting H3 antibody dynamics patterns for the H3N8 X H4N5 group that showed a boost in H3 titer (Fig 8) consistent with original antigenic sin (OAS); H4 antibodies were also detected in this challenge group (Fig 9). This phenomenon of interference was first described after sequential influenza reinfections in humans (51, 52). Older individuals can have a broader immunity through repeated exposure with highest titers to the strains individuals were exposed to early in life that are “senior strains”, a singularity known as antigenic seniority or imprinting (40, 53). Currently, we report OAS in avian hosts and between different HA subtypes (H3/H4 and possibly for H3/H10), however we expect that phenomenon could also arise between other IAV subtypes. Thus the order in which individuals are challenged with a specific virus influences the future recognition of viruses and the specificity of the responses that ultimately shapes the outcome of later exposures in life. Since population immunity influences the emergence and spread of new strains and can influence vaccination success, it is critical to have a better understanding of these processes in different host species. We need to increase our understanding of cross-reactivity patterns and boosting dynamics in re-infections to ultimately predict risks of IAV spread into different host populations in a context of non-naïve populations for instance using antibody landscapes (54).

The high degree of heterosubtypic immunity and subsequent competition found between common HA subtypes from ducks indicates that perpetuation of any subtype is dependent on the other viruses in the population and may explain the cyclic or chaotic nature in the prevalence of some subtypes. Partial immunity or complete immunity induced by pre-infection may reduce transmission potential in subsequent infections but also may promote the high degree of IAV antigenic diversity observed in wild avian populations. This competition may also result in subtype succession over time, like the predominance of H3 and H4 in fall migration (4–6) and spring blooming of other subtypes within Group 2 such as H7 or H10 (55). With equal strength of HA immunity to all subtypes the present antigenic diversity found in wild birds would be unlikely. Surprisingly, the results from H13 and H16 experimental infections in black-headed gulls showed limited cross-immunity between subtypes and suggest independent cycles for these viruses (21).

However, for some strains or subtypes, cross-immunity may not be the only factor explaining the dynamics in the pathogen population. We believe that virus fitness linked to host-specificity and functionality of IAV proteins (56) is also playing an important role as evidenced by the fact that the H12N5 IAV used in this study did not successfully replicate in mallards even though it was isolated from that host.

A possible interpretation of these results is that population immunity can reduce the probability of transmission and potential introduction success of exotic antigenic variants (57). Previous studies have reported protection to HPIAV induced by pre-exposure to LPIAV strains in different bird species (10, 13). In the context of H5N8 HPIAV clade 2.3.4.4 or other HPIAV context the present results suggest that cross-immunity could also reduce viral shedding and contribute to stopping the spread of specific virus in wild ducks that have naturally been exposed to LPIAV (58–60).

We believe that the competitive processes described here and in other studies occur in nature although in natural host populations the complexity of the system increases due to the extensive subtype diversity of co-circulating viruses that these birds are continuously exposed to. Thus, based on our results we propose that the extent of competition at individual level through host immunity could be determined by many different interacting parameters: the strains involved in infections, the exposure history (or boost responses like OAS) and likely time between exposure/s (assuming that immune memory decreases over time). These are many of the same factors that are considered in evaluating immune responses and protection against influenza in humans and domestic animals.

## Material and Methods

### Virus selection, stock preparation and titrations

All LPIAV viruses used in these trials represented common North American subtypes and were originally isolated from wild mallards in Minnesota, USA (5). Viruses included: A/Mallard/MN/AI07-4724/2007 (H3N8), A/Mallard/MN/AI11-4213/2011(H4N5), A/Mallard/MN/AI11-4982/2011(H6N2), A/Mallard/MN/AI11-4412/2011(H10N7) and A/Mallard/MN/AI11-3866/2011(H12N5). Virus stocks were propagated by a second passage in specific pathogen free (SPF) embryonated chicken eggs. Stocks were endpoint titrated using the method of Reed and Muench Method (61) in eggs to determine the median egg infectious dose (EID_50_/0.1 ml).

Ducks were inoculated with 0.1ml at an approximate dose/dilution of 10^6.0^ EID_50_/0.1ml; the inoculum was split between intranasal and oropharyngeal routes. Based on back titration, the titers of the inoculums, expressed in EID_50_/ 0.1 ml, were 10^5.8^ for H3N8 pre-challenge, 10^6.5^ for H12N8 and H6N2, 10^6.4^ for H3N8 (5 week challenge), 10^6.8^ for H10N7, 10^6.0^ for H3N8 (11 week challenge), 10^6.2^ for H4N5, 10^6.3^ for H3N8 (15 week challenge).

The HA and NA segments of the isolates were amplified (62) and later sequenced using Sanger method. Geneious (version mac6_4_8_0_4) was used for sequence alignments and to calculate amino acid distances. Sequences are publically available in GenBank (KX814369-KX814375).

### Experimental infections

All protocols for raising, infecting, and sampling mallards were approved by the IACUC at the University of Georgia (UGA; AUP#: A2013 05-021-Y1-A1).

One-day-old mallards (Murray McMurray Hatchery, Webster City, IA, USA) were raised in captivity at the animal resources facilities of the College of Veterinary Medicine, UGA. Food and water were supplied ad libitum. All individuals were identified with bands and unique ID numbers. Individuals did not exhibit behavioral changes or overt symptoms of disease during the duration of the trial. All birds gained weight before and throughout the challenge studies. At four weeks of age, 40 ducks were moved to Biosafety Level (BSL) 2+ facilities and were pre-challenged with the H3N8 LPIAV.

The study design included re-challenges with homologous and heterologous viruses (Table S1). Prior to the re-challenges, ducks were randomly divided into groups (five individuals per group, approximate ratio 1:1 females: males) and were moved into BSL 2+ poultry isolators at the Poultry Diagnostic Research Center, Athens, GA, USA that are intended to house up to five adult size mallards. After 2 days of acclimation, the re-challenge was conducted as previously described (11). Each pre-challenged group had a control group (i.e. age matched naïve individuals). Ducks were humanely euthanized at the end of all challenge trials at 14 dpi following protocols approved at the UGA. For the homologous challenge one group of five ducks was challenged with the same H3N8 at five weeks after the initial H3N8 pre-challenge and two additional groups were re-challenged with the H3N8 virus at weeks 11 and 15 post-H3N8 inoculation. Groups in the heterologous challenges were inoculated five weeks after initial H3N8 infection with subtypes representing different levels of relatedness between the key antigenic proteins HA and NA of the H3N8 used in the pre-challenge: H4N5, H10N7, H6N2 and H12N5.

### Sampling, Extraction and Virus isolations

Swabs were collected from the cloaca (CL) and oropharynx, (OP) and were placed in separate tubes containing 2 ml BHI media supplemented with antimicrobials(11). Swab samples were collected on 0-8 and on 10, 12 and 14 and 21 (only after H3N8 pre-challenge) days post infection (dpi) and kept cold until transfer to the laboratory where they were stored at −80 C° until processing.

Virus isolations was performed on swabs from all birds during all challenges on 0, 2, 4, 6, 8, 10, 12, 14 dpi as previously described (63) using two 9 to 11 days-old SPF embryonated chicken eggs. To insure that all birds were negative before any bird movement or subsequent challenge, all birds were tested by virus isolation on 21 dpi after the H3N8 pre-challenge and on 0 dpi prior to all subsequent viral challenges.

### RNA extraction and RRT-PCR

RNA from samples collected on 1 to 14 dpi were extracted using the MagMAX-96 AI/ND Viral RNA Isolation kit (Ambion, Austin, TX, USA) on the Thermo Electron KingFisher magnetic particle processor (Thermo Electron Corporation, Waltham, MA, USA) (63). Negative (BHI media) and positive (diluted isolate) controls were included in each extraction. Molecular detection of the IAV matrix gene by Real-time Reverse Transcriptase PCR (RRT-PCR) (64) was conducted with the Qiagen OneStep RT-PCR kit (Qiagen, Valencia, CA, USA) and the Cepheid SmartCycler System (Cepheid, Sunnyvale, CA, USA) (63). Negative and positive controls were used on the extraction step and an IAV matrix gene transcript (National Veterinary Services Laboratory, Ames, IA, USA) was included as positive control in the RRT-PCR. Samples with cycle threshold (Ct) values lower than 40 were considered positive and Ct–values were used as a proxy for viral load.

### Serology

Serum samples were collected from the right jugular vein and transferred into serum separator tubes (Becton, Dickinson and Company, Franklin Lakes, NJ, USA), centrifuged upon arrival to the lab and serum was stored at −20 C° until analysis. Serum samples were taken immediately prior to H3N8 pre-challenge and prior to all subsequent challenges as well as at 14 dpi and between challenges for the homosubtypic long-term groups. Serum samples were tested with a commercially available nucleoprotein (NP)-ELISA kit (bELISA; FlockChek AI MultiS-Screen antibody test kit; IDEXX Laboratories, Westbrook, Maine, USA). Specific antibodies against the different HA subtypes used in the trial were detected using a virus microneutralization (MN) assay in Madin Darby Canine Kidney cells (MDCK; ATCC, Manassas, VA, USA) as described previously (57). Sera were additionally tested using the same viruses used for inoculation in the challenge and with prototype strains for H1-H12 and H14-H15 (65) (S1 Appendix) to detect cross-reactivity. These antigens also were prepared in MDCK and tests were run using an antigen concentration of 100 TCID_50_ /25 μl.

### Statistical analysis

All analyses were run on the R software (66) using the GAMM, nlme and lme4 packages. Model selection was done using Akaike Information Criterion (67) corrected for small sample size (AICc) within the package AICcmodavg. To evaluate the viral shedding or load we used the Ct-values from the matrix RRT-PCR runs that are proportional to the RNA copy numbers from original samples. First, we studied the variation in Ct-values in pre-challenged and naïve controls for the different treatment groups with different time periods between infections. The strategy was to use linear models and include individuals as random effect using mixed models (package GAMM) due to repeated sampling of the same individuals over the course of infection. The models that were evaluated included the factors: dpi, treatment group, both factors and the interaction. Ct-values from 1 dpi were not included in the analysis as they likely residual inoculum rather than true virus replication (at 1 dpi the Ct- values were close to 40). Models including the random effect of individual and the additive effect of dpi were tested but the increased complexity of the models was penalized based on the AICc and some of them had convergence problems. In the same way, the variation of Ct-values from all H3N8 control groups including age as factor was assessed. For the heterologous re-challenge, the variation in Ct-values from control and pre-challenged groups for each of the different viruses was evaluated as described before. The total duration of infection or shedding was estimated by counting the days between inoculation and last positive virus isolations in CL swabs (which also includes cases of intermittent shedding). Birds that died or were euthanized before 14 dpi were not included in analysis. Duration of infection between groups was compared using Krustal-Wallis test.

To explore the relation between HA similarity and the degree of protection we estimated the amino acid distance between the H3 and the different HA of re-challenges and the reduction in the duration of infection within group (subtracting the mean duration of infection in pre-challenge groups to the duration in controls, calculated as the mean duration in controls minus the duration in pre-challenged as a %). The aminoacid distance was then correlated with the reduction in shedding per group.

Next, to explore the influence of different variables on isolation success (binomial response: successful or unsuccessful) we used Generalized Linear Mixed Models (GLMM) as in (49). The explanatory variables included in the models were: Ct-value, sample type (OP and CL swabs), dpi and treatment (pre-challenged or naïve individuals which means samples from a primary or secondary infection). Additionally, since individual birds were re-sampled and samples are not independent we added the individual as a random effect in the models. Interactions were not included to avoid convergence problems.

Last, MN antibody titers (as log2 transformed) between groups at a single day of sampling were compared using the Krustal-Wallis test and the paired t-test was used to compare values in two different days within groups. Samples with a titer lower than 20 were arbitrarily given a titer of log_2_ 2.5 for the model testing. The variation in antibody titers to H3 by MN was explored based on the different sampling occasions (dpi) and groups also using mixed models in the package GAMM.

One duck died (12 dpi in group H6N2) and swabs were already IAV negative, necropsy showed no gross lesions caused by LPIAV. Another duck was euthanized due difficulties walking associated to husbandry in captivity (6 dpi in group H3N8 X H6N8). These birds were not included in the analysis. The H3N8 x H12N5 group was not included in the analysis because of poor replication of this virus in the pre-exposed and naïve groups.

## Acknowledgements

We thank Morgan J. Slusher for assistance with sampling and animal care, as well to the BSL 2+ animal facilities at PDRC and Animal Resources College of Veterinary Medicine, UGA. Clara Kienzle and Nick Davis-Fields for technical assistance in the laboratory. Alexis Avril for discussions and statistical advice. NLM was funded by International Postdoc Grants from The Wenner-Gren Foundations, Stockholm, Sweden and the Swedish Research Council, VR. The research was funded by CEIRS, NIH, US under the contract HHSN272201400006C. The funding agencies were not involved in the design, implementation, or publishing of this study and the research presented herein represents the opinions of the authors, but not necessarily the opinions of the funding agencies.

---

**S1 Fig.**
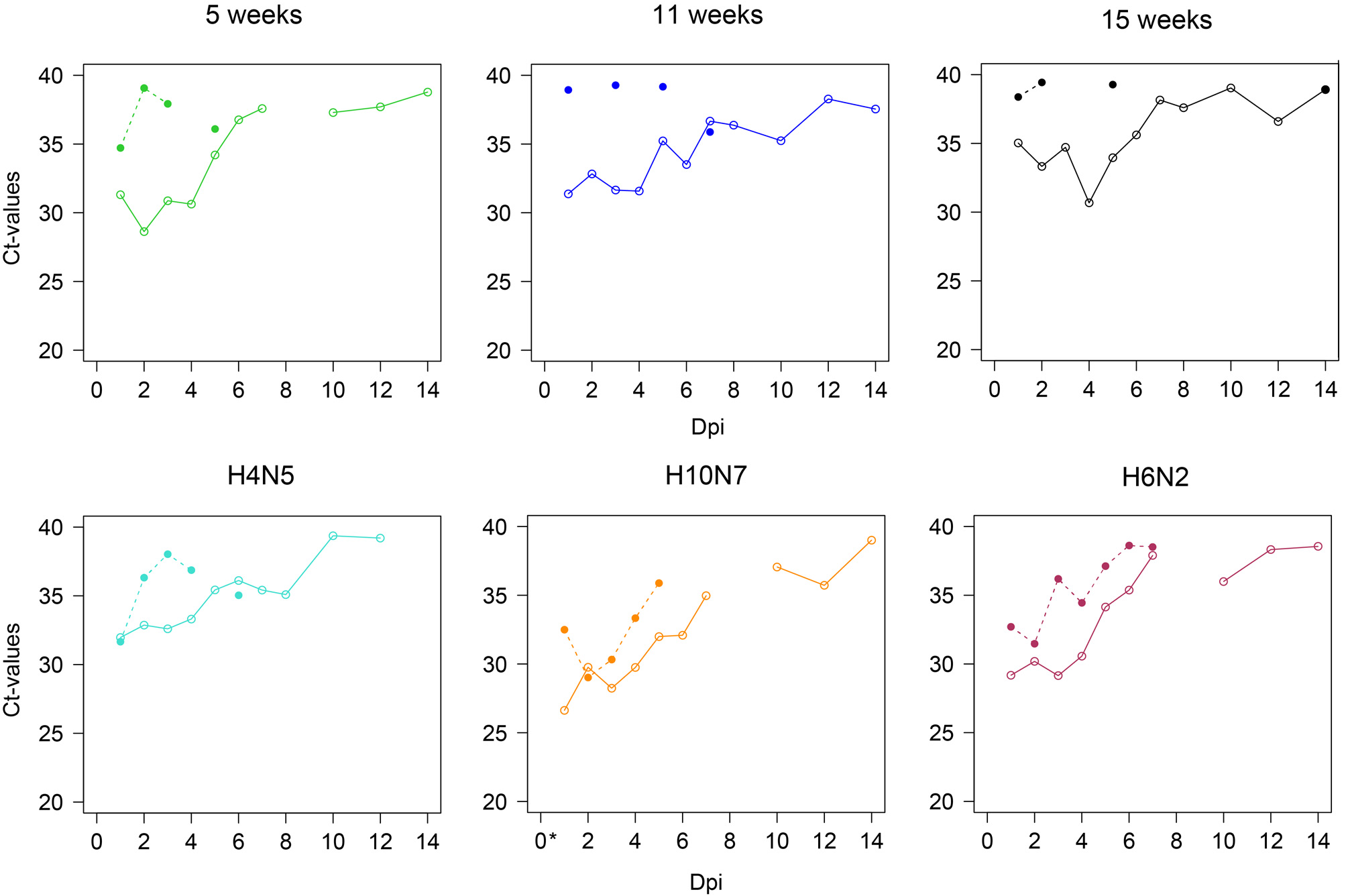
Oropharyngeal shedding showing the variation in Ct-values (average per group) for the different groups. Homologous challenge groups at infected at different time intervals are shown in top panels. Heterologous challenge groups are in lower panels). Continuous lines indicate the controls groups and discontinuous lines indicate the H3N8 pre-challenged groups (same colors as previous). * A single positive isolate from OP swab was an H10N7 detected at 0 dpi in the H10N7 control group. This duck was the last of a total of five to be inoculated in that isolator and this isolation was attributed to contamination with the inoculum. No birds had been exposed to H10N7 prior to this day.

**S2 Fig.**
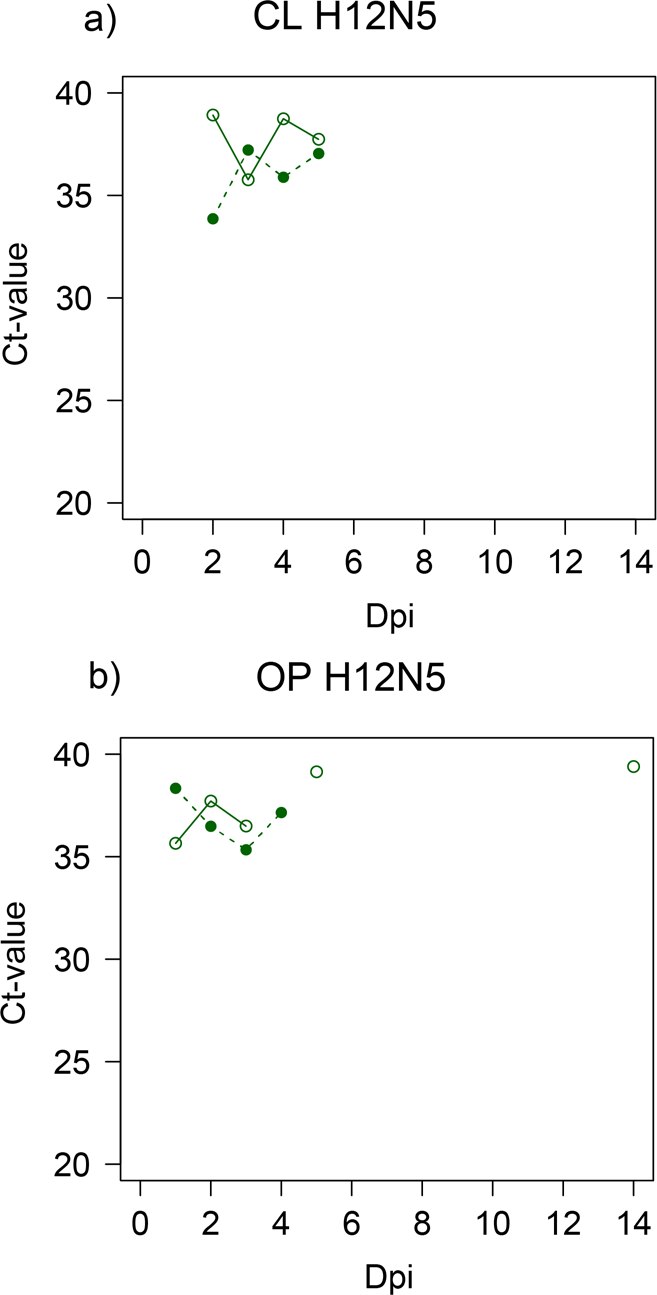
Variation in H12N5 shedding and Ct-values from 0 to 14 dpi. There was limited replication and shedding of the virus on the individuals inoculated with the H12N5. Only five CL swabs were positive by virus isolation while eight of the oropharyngeal swabs were positive by virus isolation. a) CL swabs and b) OP swabs

## References

1. Gupta S, Ferguson N, Anderson R. Chaos, persistence, and evolution of strain structure in antigenically diverse infectious agents. Science. 1998;280(5365):912–5.

2. Palese P, Wang TT. Why Do Influenza Virus Subtypes Die Out? A Hypothesis. mBio. 2011;2(5): e00150

3. Olsen B, Munster VJ, Wallensten A, Waldenstrom J, Osterhaus ADME, Fouchier RAM. Global patterns of influenza A virus in wild birds. Science. 2006;312(5772):384–8.

4. Latorre-Margalef N, Tolf C, Grosbois V, Avril A, Bengtsson D, Wille M, et al. Long-term variation in influenza A virus prevalence and subtype diversity in migratory mallards in northern Europe. P Roy Soc B-Biol Sci. 2014;281(1781).

5. Wilcox BR, Knutsen GA, Berdeen J, Goekjian V, Poulson R, Goyal S, et al. Influenza-A viruses in ducks in Northwestern Minnesota: fine scale spatial and temporal variation in prevalence and subtype diversity. PLoS ONE. 2011;6(9):e24010.

6. Munster VJ, Baas C, Lexmond P, Waldenstrom J, Wallensten A, Fransson T, et al. Spatial, temporal, and species variation in prevalence of influenza A viruses in wild migratory birds. Plos Pathog. 2007;3(5):e61. doi:10.1371/journal.ppat.0030061.

7. Hatchette TF, Walker D, Johnson C, Baker A, Pryor SP, Webster RG. Influenza A viruses in feral Canadian ducks: extensive reassortment in nature. J Gen Virol. 2004;85:2327–37.

8. Latorre-Margalef N, Grosbois V, Wahlgren J, Munster VJ, Tolf C, Fouchier RAM, et al. Heterosubtypic immunity to influenza a virus infections in mallards may explain existence of multiple virus subtypes. Plos Pathog. 2013;9(6):e1003443.

9. Jourdain E, Gunnarsson G, Wahlgren J, Latorre-Margalef N, Brojer C, Sahlin S, et al. Influenza virus in a natural host, the Mallard: experimental infection data. PLoS ONE. 2010;5(1):e8935. doi: 10.1371/journal.pone.0008935.

10. Costa TP, Brown JD, Howerth EW, Stallknecht DE, Swayne DE. Homo- and heterosubtypic low pathogenic avian influenza exposure on H5N1 highly pathogenic avian influenza virus infection in Wood Ducks (Aix sponsa). PLoS ONE. 2011;6(1):e15987. doi:10.1371/journal.pone.0015987.

11. Costa TP, Brown JD, Howerth EW, Stallknecht DE. Effect of a prior exposure to a low pathogenic avian influenza virus in the outcome of a heterosubtypic low pathogenic avian influenza infection in Mallards (Anas platyrhynchos). Avian Dis. 2010;54(4):1286–91.

12. Pepin KM, VanDalen KK, Mooers NL, Ellis JW, Sullivan HJ, Root JJ, et al. Quantification of heterosubtypic immunity between avian influenza subtypes H3N8 and H4N6 in multiple avian host species. J Gen Virol. 2012;93:2575–83.

13. Berhane Y, Leith M, Embury-Hyatt C, Neufeld J, Babiuk S, Hisanaga T, et al. Studying possible cross-protection of Canada Geese preexposed to North American low pathogenicity avian influenza virus strains (H3N8, H4N6, and H5N2) against an H5N1 highly pathogenic avian influenza challenge. Avian Dis. 2010;54(1):548–54.

14. Fereidouni SR, Starick E, Beer M, Wilking H, Kalthoff D, Grund C, et al. Highly pathogenic avian influenza virus infection of Mallards with homo- and heterosubtypic immunity induced by low pathogenic avian influenza viruses. PLoS ONE. 2009;4(8):e6706. doi: 10.1371/journal.pone.0006706.

15. Lebarbenchon C, Sreevatsan S, Lefèvre T, Yang M, Ramakrishnan MA, Brown JD, et al. Reassortant influenza A viruses in wild duck populations: effects on viral shedding and persistence in water. Proc R Soc B-Biol Sci. 2012;279(1744):3967–75.

16. Brown JD, Goekjian G, Poulson R, Valeika S, Stallknecht DE. Avian influenza virus in water: Infectivity is dependent on pH, salinity and temperature. Vet Microbiol. 2009;136(1-2):20–6.

17. Handel A, Brown J, Stallknecht D, Rohani P. A Multi-scale Analysis of Influenza A Virus Fitness Trade-offs due to Temperature-dependent Virus Persistence. PLoS Comput Biol. 2013;9(3):e1002989.

18. Kida H, Yanagawa R, Matsuoka Y. Duck influenza lacking evidence of disease signs and immune response. Infection and immunity. 1980;30(2):547–53.

19. Globig A, Fereidouni SR, Harder TC, Grund C, Beer M, Mettenleiter TC, et al. Consecutive natural influenza A virus infections in sentinel Mallards in the evident absence of subtype-specific hemagglutination inhibiting antibodies. Transboundary and Emerging Diseases. 2012:no-no.

20. Magor KE. Immunoglobulin genetics and antibody responses to influenza in ducks. Dev Comp Immunol. 2011;35(9):1008–17.

21. Verhagen JH, Hofle U, van Amerongen G, van de Bildt M, Majoor F, Fouchier RA, et al. Long-term effect of serial infections with H13 and H16 low pathogenic avian influenza viruses in black-headed gulls. J Virol. 2015.

22. Chaise C, Lalmanach AC, Marty H, Soubies SM, Croville G, Loupias J, et al. Protection Patterns in Duck and Chicken after Homo- or Hetero-Subtypic Reinfections with H5 and H7 Low Pathogenicity Avian Influenza Viruses: A Comparative Study. Plos One. 2014;9(8).

23. Fereidouni SR, Grund C, Haeuslaigner R, Lange E, Wilking H, Harder TC, et al. Dynamics of specific antibody responses induced in Mallards after infection by or immunization with low pathogenicity avian influenza viruses. Avian Dis. 2010;54(1):79–85.

24. Curran JM, Robertson ID, Ellis TM, Selleck PW, O’Dea MA. Variation in the Responses of Wild Species of Duck, Gull, and Wader to Inoculation with a Wild-Bird-Origin H6N2 Low Pathogenicity Avian Influenza Virus. Avian Dis. 2013;57(3):581–6.

25. Furger M, Hoop R, Steinmetz H, Eulenberger U, Hatt JM. Humoral Immune Response to Avian Influenza Vaccination Over a Six-Month Period in Different Species of Captive Wild Birds. Avian Dis. 2008;52(2):222–8.

26. Costa TP, Brown JD, Howerth EW, Stallknecht DE. The effect of age on avian influenza viral shedding in Mallards (Anas platyrhynchos). Avian Dis. 2010;54(1):581–5.

27. Tolf C, Latorre-Margalef N, Wille M, Bengtsson D, Gunnarsson G, Grosbois V, et al. Individual variation in influenza A virus infection histories and long-term immune responses in mallards. Plos One. 2013;8(4):e61201

28. Ferreira HL, Vangeluwe D, Van Borm S, Poncin O, Dumont N, Ozhelvaci O, et al. Differential Viral Fitness Between H1N1 and H3N8 Avian Influenza Viruses Isolated from Mallards (Anas platyrhynchos). Avian Dis. 2015;59(4):498–507.

29. Latorre-Margalef N, Gunnarsson G, Munster VJ, Fouchier RAM, Osterhaus ADME, Elmberg J, et al. Effects of influenza A virus infection on migrating mallard ducks. P Roy Soc B-Biol Sci. 2009;276(1659):1029–36.

30. Avril A, Grosbois V, Latorre-Margalef N, Gaidet N, Tolf C, Olsen B, et al. Capturing individual-level parameters of influenza A virus dynamics in wild ducks using multistate models. Journal of Applied Ecology. 2016: 1365–2664.

31. Grebe KM, Yewdell JW, Bennink JR. Heterosubtypic immunity to influenza A virus: where do we stand? Microbes Infect. 2008;10(9):1024–9.

32. Kreijtz JHCM, Fouchier RAM, Rimmelzwaan GF. Immune responses to influenza virus infection. Virus Res. 2011;162(1-2):19–30.

33. Seo SH, Webster RG. Cross-reactive, cell-mediated immunity and protection of chickens from lethal H5N1 influenza virus infection in Hong Kong poultry markets. J Virol. 2001;75(6):2516–25.

34. Corti D, Voss J, Gamblin SJ, Codoni G, Macagno A, Jarrossay D, et al. A neutralizing antibody selected from plasma cells that binds to group 1 and group 2 influenza A hemagglutinins. Science. 2011;333(6044):850–6.

35. Ekiert DC, Friesen RHE, Bhabha G, Kwaks T, Jongeneelen M, Yu WL, et al. A highly conserved neutralizing epitope on group 2 influenza A viruses. Science. 2011;333(6044):843–50.

36. Sui J, Sheehan J, Hwang WC, Bankston LA, Burchett SK, Huang CY, et al. Wide prevalence of heterosubtypic broadly neutralizing human anti-influenza A antibodies. Clin Infect Dis. 2011;52(8):1003–9.

37. Wang TT, Palese P. Universal epitopes of influenza virus hemagglutinins? Nat Struct Mol Biol. 2009;16(3):233–4.

38. Pica N, Hai R, Krammer F, Wang TT, Maamary J, Eggink D, et al. Hemagglutinin stalk antibodies elicited by the 2009 pandemic influenza virus as a mechanism for the extinction of seasonal H1N1 viruses. P Natl Acad Sci USA. 2012;109(7):2573–8.

39. Wohlbold TJ, Krammer F. In the Shadow of Hemagglutinin: A Growing Interest in Influenza Viral Neuraminidase and Its Role as a Vaccine Antigen. Viruses-Basel. 2014;6(6):2465–94.

40. Worobey M, Han GZ, Rambaut A. Genesis and pathogenesis of the 1918 pandemic H1N1 influenza A virus. P Natl Acad Sci USA. 2014;111(22):8107–12.

41. Gostic KM, Ambrose MR, Worobey M, Lloyd-Smith JO. Potent Protection against H5N1 and H7N9 Influenza via Childhood Hemagglutinin Imprinting. bioRxiv. 2016.

42. Bahl J, Vijaykrishna D, Holmes EC, Smith GJD, Guan Y. Gene flow and competitive exclusion of avian influenza A virus in natural reservoir hosts. Virology. 2009;390(2):289–97.

43. Bahl J, Krauss S, Kuhnert D, Fourment M, Raven G, Pryor SP, et al. Influenza A Virus Migration and Persistence in North American Wild Birds. Plos Pathog. 2013;9(8).

44. Katzelnick LC, Fonville JM, Gromowski GD, Arriaga JB, Green A, James SL, et al. Dengue viruses cluster antigenically but not as discrete serotypes. Science. 2015;349(6254):1338–43.

45. Read AF, Baigent SJ, Powers C, Kgosana LB, Blackwell L, Smith LP, et al. Imperfect Vaccination Can Enhance the Transmission of Highly Virulent Pathogens. PLoS Biol. 2015;13(7):e1002198.

46. Cattoli G, Fusaro A, Monne I, Coven F, Joannis T, El-Hamid HSA, et al. Evidence for differing evolutionary dynamics of A/H5N1 viruses among countries applying or not applying avian influenza vaccination in poultry. Vaccine. 2011;29(50):9368–75.

47. Koel BF, van der Vliet S, Burke DF, Bestebroer TM, Bharoto EE, Yasa IWW, et al. Antigenic Variation of Clade 2.1 H5N1 Virus Is Determined by a Few Amino Acid Substitutions Immediately Adjacent to the Receptor Binding Site. mBio. 2014;5(3).

48. Koel BF, Burke DF, Bestebroer TM, van der Vliet S, Zondag GCM, Vervaet G, et al. Substitutions Near the Receptor Binding Site Determine Major Antigenic Change During Influenza Virus Evolution. Science. 2013;342(6161):976–9.

49. Latorre-Margalef N, Avril A, Tolf C, Olsen B, Waldenstrom J. How Does Sampling Methodology Influence Molecular Detection and Isolation Success in Influenza A Virus Field Studies? Appl Environ Microb. 2016;82(4):1147–53.

50. Brown JD, Poulson R, Carter DL, Lebarbenchon C, Stallknecht DE. Infectivity of avian influenza virus-Positive field aamples for mallards: What do our diagnostic results mean? Journal of wildlife diseases. 2013;49(1):180–5.

51. Francis T. Influenza - the Newe Acquayantance. Ann Intern Med. 1953;39(2):203–21.

52. Webster RG. Original Antigenic Sin in Ferrets - Response to Sequential Infections with Influenza Viruses. J Immunol. 1966;97(2):177-&.

53. Kucharski AJ, Lessler J, Read JM, Zhu HC, Jiang CQ, Guan Y, et al. Estimating the Life Course of Influenza A (H3N2) Antibody Responses from Cross-Sectional Data. Plos Biology. 2015;13(3).

54. Fonville JM, Wilks SH, James SL, Fox A, Ventresca M, Aban M, et al. Antibody landscapes after influenza virus infection or vaccination. Science. 2014;346(6212):996–1000.

55. Ramey AM, Poulson RL, Gonzalez-Reiche AS, Wilcox BR, Walther P, Link P, et al. Evidence for Seasonal Patterns in the Relative Abundance of Avian Influenza Virus Subtypes in Blue-Winged Teal (Anas discors). Journal of wildlife diseases. 2014;50(4):916–22.

56. Galloway SE, Reed ML, Russell CJ, Steinhauer DA. Influenza HA Subtypes Demonstrate Divergent Phenotypes for Cleavage Activation and pH of Fusion: Implications for Host Range and Adaptation. Plos Pathog. 2013;9(2).

57. Ramey AM, Poulson RL, Gonzalez-Reiche AS, Perez DR, Stallknecht DE, Brown JD. Genomic Characterization of H14 Subtype Influenza A Viruses in New World Waterfowl and Experimental Infectivity in Mallards (Anas platyrhynchos). Plos One. 2014;9(5).

58. Verhagen JH, van der Jeugd HP, Nolet BA, Slaterus R, Kharitonov SP, de Vries PP, et al. Wild bird surveillance around outbreaks of highly pathogenic avian influenza A(H5N8) virus in the Netherlands, 2014, within the context of global flyways. Eurosurveillance. 2015;20(12).

59. Krauss S, Stallknecht DE, Slemons RD, Bowman AS, Poulson RL, Nolting JM, et al. The enigma of the apparent disappearance of Eurasian highly pathogenic H5 clade 2.3.4.4 influenza A viruses in North American waterfowl. Proc Natl Acad Sci U S A. 2016;113(32):9033–8.

60. Lee DH, Torchetti MK, Winker K, Ip HS, Song CS, Swayne DE. Intercontinental Spread of Asian-Origin H5N8 to North America through Beringia by Migratory Birds. J Virol. 2015;89(12):6521–4.

61. Reed LJ, Muench H. A simple method of estimating fifty per cent endpoints Am J Epidemiol. 1938;27(3):493–7.

62. Bragstad K, Jorgensen PH, Handberg KJ, Mellergaard S, Corbet S, Fomsgaard A. New avian influenza A virus subtype combination H5N7 identified in Danish mallard ducks. Virus Res. 2005;109(2):181–90.

63. Brown JD, Berghaus RD, Costa TP, Poulson R, Carter DL, Lebarbenchon C, et al. Intestinal excretion of a wild bird-origin H3N8 low pathogenic avian influenza virus in mallards (Anas Platyrhynchos). Journal of wildlife diseases. 2012;48(4):991–8.

64. Spackman E, Senne DA, Myers TJ, Bulaga LL, Garber LP, Perdue ML, et al. Development of a real-time reverse transcriptase PCR assay for type A influenza virus and the avian H5 and H7 hemagglutinin subtypes. Journal of clinical microbiology. 2002;40(9):3256–60.

65. Latorre-Margalef N, Ramey AM, Fojtik A, Stallknecht DE. Serologic Evidence of Influenza A (H14) Virus Introduction into North America. Emerg Infect Dis. 2015;21(12):2257–9.

66. Team RC. R: A language and environment for statistical computing. Vienna, Austria: R Foundation for Statistical Computing; R version 2.14.0, 2011.

67. Kenneth P. Burnham DRA. Model selection and multimodel inference: A practical information-theoretic approach. 2nd ed: Springer Science+ Bussiness Media, Inc.; 2002.

